# Thermal fluctuations independently modulate physiological plasticity and the dynamics of the gut microbiome in a tropical rocky shore oyster

**DOI:** 10.1101/2023.02.08.527599

**Authors:** Bovern Suchart Arromrak, Adrian Tsz Chun Wong, Tin Yan Hui, Kin Sum Leung, Gray A. Williams, Monthon Ganmanee, Thierry Durand, Jetty Chung Yung Lee, Juan D. Gaitan-Espitia

## Abstract

Extreme high thermal conditions on tropical rocky shores are challenging to the survival of intertidal ectotherms. Yet, many species are highly successful in these environments in part due to their ability to regulate intrinsic mechanisms associated with physiological stress and their metabolic demand. More recently, there has been a growing awareness that other extrinsic mechanisms, such as animal-associated microbial communities, can also influence the tolerance and survival of ectotherms under stressful conditions. However, the extent to which the intrinsic and extrinsic mechanisms are functionally linked as part of the overall adaptive response of intertidal animals to temperature change and stress is poorly understood. Here, we examined the dynamics and potential interactions of intrinsic and extrinsic mechanisms in the tropical high-supratidal oyster, *Isognomon nucleus*. We found that oysters modulate their internal biochemistry (oxidized PUFA products, including 5-F_2t_-IsoP, 10-F_4t_-NeuroP, 13-F_4t_-NeuroP, and 16-F_1t_-PhytoP) as part of their adaptive regulation to cope with physiological stress during periods of extreme high temperatures when emersed. However, while we detected variation in alpha diversity (ASV richness and Shannon diversity index), dominant microbial taxa and microbial functions across time, no association was found with the host biochemical profiles. The findings here suggest that the thermal condition within oysters can independently influence their intrinsic biochemical responses and extrinsic microbiome profiles. Together, these mechanisms may contribute to the thermal tolerance and survival of the oysters in the challenging conditions of the tropical high-supratidal zone.

## Introduction

The tropical rocky shore is one of the most extreme environments on Earth. This system is characterized by a marked heterogeneous thermal landscape influenced by tidal dynamics (Raffaelli and Hawkins, 1999; Helmuth and Hofmann, 2001). Across this vertical gradient defined by the tides, environmental temperatures shape the physiology, behaviour, and spatial distribution of tropical intertidal ectotherms (Garrity, 1984; Williams and Morritt, 1995; Chow, 2004; Samakraman *et al*., 2010; Giomi *et al*., 2016; Ng *et al*., 2017). For ectotherms occupying the high intertidal zone (hereafter, high-supratidal zone), thermal and oxidative stress (i.e., excessive production of reactive oxygen species - ROS) during desiccation periods of extended emersion, are the two of the major physiological challenges faced (Sokolova and Pörtner, 2001; Freire *et al*., 2011; Birben *et al*., 2012).

To survive under these challenging conditions, high-supratidal ectotherms modulate intrinsic/innate mechanisms associated with their internal physiology (e.g., metabolic rate) and biochemical activities. This modulation can be observed through, for instance, the removal of ROS via polyunsaturated fatty acids (PUFA; Sokolova and Pörtner, 2001; Yin *et al*., 2017; Monroig *et al*., 2022), and/or the control of the gene expression associated with heat shock response (HSR) and oxidative stress response (OSR; Wang *et al*., 2022). PUFAs such as Arachidonic acid (ARA) and <u>D</u>ocosa<u>h</u>exaenoic <u>a</u>cid (DHA) are known to be effective molecules and agent for ROS removal, because they contain unsaturated double bonds making them susceptible to ROS reaction (particularly to peroxide, HO•) (Sargent, 1976; Leung *et al*., 2015; Yin *et al*., 2017; Fadhlaoui and Lavoie, 2021). The non-enzymatic peroxidation of these PUFAs, generates by-products that can be used as biomarkers of the ROS level (Miller *et al*., 2014; Yonny *et al*., 2016; Galano *et al*., 2017). In addition to the biochemical modulation mechanism, some intertidal organisms (e.g., the oysters, *Isognomon nucleus*) are able to enter into a hypometabolic state (i.e., metabolic depression, for instance, marked by a drop in heart rate; Marshall and McQuaid, 2011; Hui *et al*., 2020) to avoid excessive energy consumption when exposed to high temperatures during low tides. Together, these functional characteristics are regarded as important adaptive traits for intertidal ectotherms to cope and ameliorate the extreme stress experienced on tropical rocky shores.

Survival under stressful environmental conditions is also hypothesized to be influenced by extrinsic/acquired mechanisms such as host-associated microbial communities (i.e., microbiome; Bang *et al*., 2018; Ahmed *et al*., 2019). The structure and functions of these microbiomes have been suggested to play important roles in shaping tolerances and biochemical regulation of animals to elevated temperatures (Herrera *et al*., 2020; Sepulveda and Moeller, 2020; Jaramillo and Castañeda, 2021). Under such conditions, functional microbiomes that are resilient/responsive to environmental changes are fundamental to the host’s health and survival (Soen, 2014; Kremer *et al*., 2018). For instance, Epstein et al. (2019) demonstrated that the resilience of the coral, *Pocillopora acuta,* to a stressful thermal event may be attributed to the stability of the microbiome composition and structure. In contrast, a dysbiotic host state is commonly associated with a destabilized microbiome (Fan *et al*., 2013; McDevitt-Irwin *et al*., 2019). This is typically reflected by a marked variation among host-associated microbiomes, a signal indicating the host losing control of the microbial community (e.g., increase in opportunistic pathogens) and/or differential sensitivities of microbiomes to extreme disturbances (e.g., elevated temperature; Carey and Duddleston, 2014; Petersen and Round, 2014; Lokmer and Wegner, 2015). This pattern is in-line with the “Anna Karenina Principle”, which states, “all happy families are all alike; each unhappy family is unhappy in its own way” (Zaneveld *et al*., 2017; Díaz-Almeyda *et al*., 2022). In other words, stressed organisms may host more stochastic microbiomes than healthy organisms. Although microbiome stability seems important for host homeostasis, some studies have suggested that flexibility in the microbiome can also provide adaptive capacities to the host, allowing it to cope with rapidly changing environments (Maher *et al*., 2020; Voolstra and Ziegler, 2020). Ziegler et al. (2017), for example, showed that the coral *Acropora hyacinthus* exhibits changes in its microbiome composition and structure that are aligned with the local thermal environmental conditions. Similarly, several studies have also shown seasonal variation in the host-associated microbiome of many marine organisms such as bivalves and echinoderms, as a potential plastic/flexible response to environmental change (Pierce and Evan, 2019; Feng *et al*., 2021). Microbiome stability or flexibility may, therefore, be a sign of host adaptive responses to environmental changes. However, the influence of these contrasting patterns is context-dependent and thus, their contribution to the tolerances and survival of animal hosts may differ in thermally challenging environments such as tropical rocky shores.

Previous studies have shown that the capacity for plastic adaptive responses of animal hosts to environmental stress may be modulated by the interactions between intrinsic (physiology and biochemical changes of the host) and extrinsic (host’s microbiome structure and function) mechanisms (Muñoz *et al*., 2019; Marangon *et al*., 2021; Fontaine and Kohl, 2023). It is unclear, however, to what extent these mechanisms are functionally linked as part of the whole organism’s adaptive response (i.e., holobiont: host and microbiome) to survive in an extreme environment such as the tropical intertidal rocky shore. In this study, we investigated the intrinsic and extrinsic mechanisms underpinning the tolerances and survival of a tropical intertidal ectotherm (oyster, *Isognomon nucleus*) by assessing the influence of natural variation in environmental temperature on i) host physiology (i.e., biochemical dynamics) ii) the diversity, structure, and function of the host gut-associated microbiome and iii) the interplay between these mechanisms. The integration of intrinsic and extrinsic mechanisms offers a holobiont perspective for understanding the stress tolerance regulation of ectotherms in extreme high temperature and fluctuating environments (Alberdi *et al*., 2016; Apprill, 2017; Bang *et al*., 2018).

## Materials and methods

### Site and species descriptions

Host-microbiome interactions were investigated in the oyster, *Isognomon nucleus* (Bivalvia: Isognomonidae), a dominant ectotherm of the high-supratidal zone in Southeast Asia. Animals were collected from Ko Sichang, Thailand (latitude: 13°08’52’’N; longitude: 100°48’11’’E). In this area, *I. nucleus* is abundant and forms a distinctive band at the high-supratidal zone (∼2.5 - 3 m above the mean sea level; Samakraman *et al*., 2010), where rock temperatures vary daily (emersion period) from ∼30°C at dawn to ∼50°C at noon (hot season: March to June; Hui *et al*., 2020). The oyster exhibits strong variations in physiological and metabolic adjustments with elevated body temperatures, providing an ideal system to investigate the functional role of the microbiome to facilitate the host’s survival in this thermally extreme and variable environment (Giomi *et al*., 2016; Hui *et al*., 2020).

### Sample collection and processing

A total of 80 individuals of *I. nucleus* were randomly selected at four different timepoints within an emersion period (Fig. 1A; 20 animals at each time). These timepoints (09:15 hr, 12:30 hr, 16:12 hr, and 17:40 hr; hereafter, morning, noon, early-afternoon, and late-afternoon, respectively), encompass a temporal cycle characterized by major changes in the environmental temperature (Table S1) as a result of the tidal dynamics and air/sun exposure (Hui *et al*., 2020). Body temperatures were measured from 10 randomly selected individuals at each timepoint using a digital thermometer (± 0.1 °C, TM-947SD, Lutron, Taiwan) where a fine-tipped type-K thermocouple was inserted into the bodies through the valve opening. At each time point, the whole tissue of five collected oysters was quickly removed at the site with a sterile dissecting kit and snap-frozen in liquid nitrogen on shore for oxidative stress quantification (as a proxy for host physiological and biochemical dynamics). The guts of another five individuals were also dissected at each timepoint on-site, snap frozen in liquid nitrogen for gut microbiome assessment. All collected samples (20 for oxidative stress and 20 for microbiome analysis across four time points) were stored in a portable liquid nitrogen tank and transported to The University of Hong Kong within three days, where they were held in a −80℃ freezer until processing and analyses. Because of the weather instability and strong waves during the fieldwork, temporal sampling was limited to one day to comply with logistic and safety regulations.

**Fig. 1.**
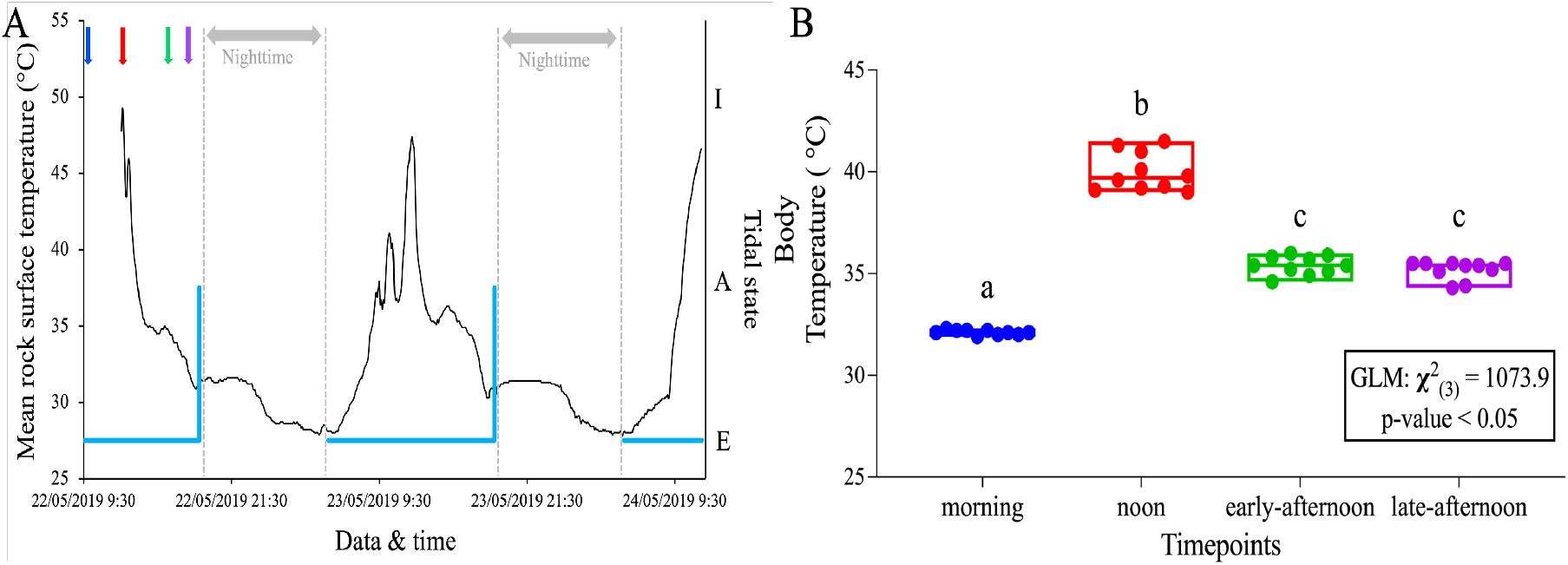
The body temperature of *Isognomon nucleus* varies across sampling timepoints within a tidal cycle. (A) Thermal (black line) and tidal conditions (blue line) as experienced by *I. nucleus* at the study site. Thermal condition is represented by the mean rock surface temperature (averaged over 3 – 5 i-Button temperature loggers), and tidal condition during the day is categorized into three states: “E” (exposed), “A” (awash), and “I” (immersed). Coloured arrows indicate the sampling time points of *I. nucleus* (colours match to boxplot in Fig. 1B) to assess its physiological responses (i.e., biochemical dynamics) and microbiome taxonomic/functional profile (B) Boxplots showing *I. nucleus* body temperatures (n=10) across different timepoints (Centre line represent the median; box limits indicate the first and third quartiles). Statistical significance (p-value < 0.05) is indicated if the letters above each bar are different.

### Rock surface temperature and tidal dynamics

To quantify environmental conditions experienced by *Isognomon nucleus* throughout the tidal cycle, rock surface temperatures and tidal levels were recorded. To measure and track surface rock temperature, i-Button loggers (DS1922L, Maxim, US) were firstly embedded in waterproofing resin (Scotchcast, 3M, US), placed in plastic Falcon tube caps and affixed, using marine epoxy (A-788, Z-Spar, US; Ng *et al*., 2021), onto the shore at the height where *I. nucleus* were collected. Rock surface temperatures were logged by these i-Buttons every 5 minutes at a resolution of 0.5 °C (n = 5). Additionally, a time-lapse camera (TLC200Pro, Brinno, Taiwan) was set adjacent to the study site to monitor the immersion state of the *I. nucleus* band during the day (∼ 05:00 hr to 18:00 hr) to assess the tidal states. Images were captured every 10 minutes and the tidal level was scored according to whether the band was underwater (immersed), exposed to air (emersed), or being splashed by waves during transitions from immersion/emersion (awash). During the emersion period, animals showed no intermittent gaping (Hui, T.Y., unpublished data). Predicted tide height from the lowest low water height on the sampling date was also obtained from the Hydrographic Department, Royal Thai Navy Website (https://www.hydro.navy.mi.th/index1.php#nogo2): Lowest tidal height – 0.55m (12:56 hr) and highest tidal height - 3.36m (20:53hr).

### Determination of oxidized polyunsaturated fatty acids products

The levels of oxidative stress (i.e., associated with ROS production) of the oysters were determined based on the concentration of non-enzymatic oxidized polyunsaturated fatty acids (PUFA) products (hereafter, oxidized PUFA products; Durand *et al*., 2009; Basu, 2010; Leung *et al*., 2015). These oxidized products are derived from phospholipids in the membranes, hence require hydrolysis for measurement. A modified Folch extraction method was used to extract the oxidized PUFA products from the oysters sampled (n=5) at each timepoint (t=4) (Folch *et al*., 1957; Dupuy *et al*., 2016). In brief, 0.05g of oyster flesh was suspended in a 5 ml ice-cold Folch solution (chloroform: methanol, 2:1, v/v) with 0.01% <u>B</u>utylated <u>H</u>ydroxy<u>T</u>oluene (BHT; used as anti-oxidant to prevent lipid peroxidation during sample processing) (w/v) and homogenized at 24,000 rpm in 20 secs bursts on ice using a polytron benchtop homogenizer (T25, Ultra-Turrax, IKA, Germany). Homogenization was performed twice, and the two parts of the Folch solution extracts were pooled. The pooled solutions were added with 2 ml of 0.9% NaCl to introduce phase separation. The mixture was then shaken on ice for 30 min and centrifuged at 2000 × g for 10 min at room temperature. The lower chloroform phase was transferred into a 30 ml glass vial. The extracted samples were then added with potassium hydroxide (1M in methanol) and 1X PBS (pH7.4). Samples were then shaken overnight in the dark at room temperature for alkaline hydrolysis of the extracted samples. The reaction was terminated by adding 1M of hydrochloric acid, methanol, 20 mM of formic acid, and 40 mM of formic acid. Deuterated internal standards (5ng prepared in methanol) were also added after the hydrolysis. Purification of the oxidized PUFA metabolites was performed with the mixed anion solid phase extraction (MAX SPE, Waters, Milford, MA USA). Before the start of the solid phase extraction, the SPE column was preconditioned with methanol and 20 mM formic acid. One sample was then loaded into a column and cleaned with 2% ammonium hydroxide and 20 mM formic acid. The final eluent was collected with hexane/ethanol/acetic acid (70/29.4/0.6, v/v/v). All the samples were then placed on a heating block and heated at 37°C under a stream of nitrogen gas until they were completely dried. The dried samples were then re-suspended in 100 µL methanol and cleaned through a 0.45 µm PTFE filter to remove insoluble impurities, and then immediately analysed by liquid chromatography-tandem mass spectrometry (LC-MS/MS).

In our study, we used the Sciex X500R QTOF LC-MS/MS system (Sciex Applied Biosystems, MA, USA) which consisted of an Exion LC liquid chromatography with a C18 column (150×2.1 mm, 2.6 µm particle size, Phenomenex, Torrance, CA, USA) which was maintained at 40°C for LC-MS/MS analysis. A mobile phase consisting of 0.1% aqueous acetic acid (solvent A) and 0.1% acetic acid/methanol (solvent B) was used. The flow rate was set to 300 µl/min and the injection volume to 10 µl. For each sample analysis, the gradient condition was maintained at 20% of solvent B for 2 min and between 20% and 98% for 8 min and then held for 5 min. Then, the percentage of solvent B was reduced to 20% in 1 min and held for an additional 5 min to equilibrate to the initial conditions. The X500R QTOF system was operated in negative electrospray ionization (ESI) mode. The spray voltage was set to −4500V and nitrogen was used as the curtain gas. The ionization chamber temperature was set at 350°C, and the pressure of the ion source gases 1 and 2 were 35 and 45 psi, respectively. The declustering potential (DP) was set to −80V and the collision energy (CE) was −10V for the QTOF MS. The scan mode was set to multiple reaction monitoring (MRM). All data collected by the X500R QTOF system was analyzed by the Sciex operating system (version 1.2.0.4122). The quantification of each analyte was determined by correlating the peak area to its corresponding deuterated internal standard peak. For analytes without corresponding deuterated internal standards, the following deuterated internal standards, 5(*S*)-HETE-d_8_, 15-F_2t_-IsoP-d_4,_ and 4-F_4t_-NeuroP-d_4_ were used for quantification. We identified fifteen oxidized PUFA products that are known to be the products of non-enzymatic lipid peroxidation (Table S2), based on previous work by Dupuy et al.(2016), Galano et al.(2017) and Lee et al.(2018).

### DNA extraction and 16S rRNA gene sequencing

Total genomic DNA was extracted from the oysters’ guts (∼0.03g) using the commercial DNA extraction kit, Dneasy Power Soil Kit (Qiagen GmbH, Hilden, Germany), following the manufacturers protocol with some modifications (i.e., the solution mixture was incubated at 55℃ in a water bath for 30 minutes after the bead-beating step). The extracted DNA (100μl) was then collected in a microcentrifuge tube and preserved in a −80℃ freezer. All the DNA samples were then sent to Ramaciotti Centre for Genomics at the University of New South Wales, Sydney Australia for 16S rRNA amplicon paired-end sequencing (2X 300bp) on an Illumina Miseq Platform. Before the amplicon sequencing, the V3-V4 variable region of the 16S rRNA gene from each DNA sample was amplified using the following primers: (1) 341F (Forward Primer);5’-CCTACGGGNGGCWGCAG-3’ (2) 805R (Reverse Primer); 5’-GACTACHVGGGTATCTAATCC-3’(Herlemann *et al*., 2011; Klindworth *et al*., 2013). Raw reads received were the demultiplexed paired-end sequences in a FASTQ format and they were deposited in NCBI Sequence Read Archive (SRA) under Bioproject number PRJNA931123. The data will be made available once published.

### 16S rRNA gene sequence processing

Raw demultiplexed paired-end sequences were processed following the Quantitative Insights into Microbial Ecology (QIIME2 ver. 2021.4) pipeline (Bolyen *et al*., 2019). Briefly, the raw data was first visualized with the demux plug-in function for quality-checking and identifying poor-quality base regions to filter. The sequences were then trimmed, filtered, denoised, chimeras-removed, and merged using the DADA2 plug-in function in QIIME2 (Callahan *et al*., 2016). The sequences were identified at a single nucleotide threshold (Amplicon Sequence Variants; ASV; Callahan *et al*., 2017). Additional steps of quality control were performed to reduce non-target sequences by aligning (vsearch alignment method) the ASV sequences to the 99% representative sequences from the greengenes database and only ASV with a similarity threshold of 0.65% and coverage threshold of 0.5% were retained following https://github.com/biocore/deblur/issues/139#issuecomment-282143373. Singletons and sequences with less than 349bp were also removed with filter-features and filter-sets function in QIIME2, respectively. The quality-filtered sequences were then taxonomically assigned with a naïve-Bayes classifier trained on the V3-V4 region of the 16S gene in the Silva v138 database (Quast *et al*., 2013), using the classify-sklearn function in the feature-classifier plugin (Bolyen *et al*., 2019). The output from this generated a taxonomy table with taxonomic naming for each ASV found within our samples. ASV sequences that were identified as mitochondria, chloroplast, archaea, and eukaryotes were filtered out. The quality-filtered ASV sequences were then ready to be used for downstream analysis.

### Profiling microbiome functions

The functional microbiome was predicted using the software Phylogenetic Investigation Communities by Reconstruction of Unobserved States version 2 (PICRUSt2; Douglas *et al*., 2020). The predicted functions associated with the microbiome (using the quality-filtered sequences) were inferred based on the MetaCyc pathway abundance, using the PICRUSt plug-in within QIIME2.

### Data analysis

Unless otherwise specified, the R statistical programming language (ver. 4.2.0, R Core Team) and R Studio was used to generate graphics and statistical analyses. The oysters’ body temperature data violated the statistical assumption of homogeneity of variances (Levene’s test, p-value < 0.05). Hence, the statistical analysis for this trait across different timepoints was assessed using a generalized linear model (GLM) with a gamma distribution. The likelihood-ratio chi-square test was used to determine the statistical significance and is indicated if the p-value was < 0.05 (R package “Car”; Fox and Weisberg, 2019). Post-hoc testing for pairwise comparison was made using the estimated marginal means with Tukey’s tests from the R package emmeans (Lenth, 2023)

Individual oxidized PUFA products were first evaluated for normality (Shapiro-Wilk’s Test) and homogeneity of variance (Levene’s test) assumptions. Metabolites that fulfilled both assumptions were tested with one way Analysis of variance (ANOVA) to investigate the variation in metabolites’ concentrations across timepoints, followed by *post-hoc* Tukey’s Honest significant difference (Tukey’s HSD). For metabolites that violated either one of the assumptions, they were evaluated with GLM (gamma distribution) following a similar procedure as with the body temperature datasets.

Assessment of oxidized PUFA products’ beta diversity (with all the 15 metabolites or only the 4 significant metabolites) was performed by first generating a Euclidean distance matrix between samples, using normalized metabolites datasets (i.e., autoscaling; van den Berg *et al*., 2006). Euclidean distance metric was selected as it allows for accurate and reliable analysis for continuous datasets and testing the differences between groups of samples based on the metabolite data (Qi and Voit, 2017; Mallick *et al*., 2019; Nguyen *et al*., 2021). A principle Coordinate Analysis (PCoA) plot was then created to visualize the oxidized PUFA products clustering across different timepoints, using the ggplot2 R package (Wickham, 2008). Permutational multivariate analysis of variance (PERMANOVA) was performed with 999 permutations to test for the differences in oxidized PUFA products profile across timepoints (Oksanen *et al*., 2022). We also performed a permutational multivariate dispersion test (PERMDISP) to assess differences in the dispersion of metabolite profiles between timepoints with 999 permutations followed by pairwise comparisons. All multivariate metabolites analyses were performed using the package vegan in R (Oksanen *et al*., 2022).

Oysters’ gut microbiome analysis was conducted by first combining the ASV count data, taxonomy table, and metadata into a simplified data matrix for ease of downstream processing, using the phyloseq() function from the phyloseq R package (McMurdie and Holmes, 2013). Rarefaction curves (total ASVs captured against sampling depth) were generated for each sample with the ggrare() function from the ranacapa R package (Kandlikar *et al*., 2018). All samples approached the asymptote level, indicating that the total ASV richness for each sample has been captured (Fig. S1). Each of the samples was then rarefied to a sampling depth of 21,282 reads (a cut-off that balances the maximum number of sequences retained per sample and minimizing the number of samples removed) for downstream analysis (Weiss *et al*., 2017; i.e., alpha and beta diversity; McKnight *et al*., 2019). Using the rarefied data, the microbiome alpha diversity (i.e., the structure of a microbial community with respect to its richness and evenness; Willis, 2019) was determined based on two indices – ASV richness (based on the number of ASV present within a group) and Shannon-Wiener (H’) diversity (based on both, the number of ASV and their abundances within a group). ASV richness was calculated with the Base-R function, while Shannon-Wiener (H’) diversity index was generated using the diversity() function from the Vegan package (Oksanen *et al*., 2022). Differences in the microbiome alpha diversity across the timepoints were assessed with ANOVA and followed by a *post-hoc* Tukey HSD test. The ANOVA statistical assumptions of normality (Shapiro-Wilk test) and homogeneity of variance (Levene’s test) for all datasets were tested and fulfilled (p-value > 0.05). Linear regression analysis was also conducted to determine the relationship between the oysters’ microbiome alpha diversity (i.e., ASV richness and Shannon-Wiener (H’) index) and body temperature (i.e., average of the 10 random independent body temperature replicates collected at each timepoint). The statistical assumptions for the linear regression model were fulfilled based on the normality test (Shapiro-Wilk test) and residuals plot (variance homogeneity).

While alpha diversity focuses on community variation within a community (sample), beta diversity quantifies (dis-)similarities between communities/samples in terms of species presence/absence and abundance (Kers and Saccenti, 2022). Metrics of beta diversity are typically used to measure species turnover or replacement across different locations, environmental conditions or timepoints (Kers and Saccenti, 2022). Here, we assessed beta diversity of the oysters’ gut microbiome to evaluate temporal differences in taxonomic and functional structure across all samples simultaneously. Bray-Curtis dissimilarity distance matrix was first generated between all samples. Then, a PCoA plot was created to visualize microbiome taxonomic and functional clusters across different timepoints, using the ggplot2 R package (Wickham, 2008). The chosen dissimilarity distance metric (Bray-Curtis) followed findings by Mcknight et al. (2019) and Weiss et al. (2017), in which authors showed optimal and high confidence for community level comparison with rarefied microbiome datasets. PERMANOVA was performed with 999 permutations to test for the differences in the microbiome composition and structure between timepoints (Oksanen *et al*., 2022). PERMDISP with 999 permutations was performed to test for differences in the taxonomic and functional microbiome profile dispersion followed by pairwise comparison. All multivariate analyses were conducted using the package vegan in R.

Differential abundance analysis of each microbial taxa was performed with family taxonomic level data (using unrarefied quality-filtered ASV sequences) and analysed with Analysis of Composition Microbiome with Bias Correction (ANCOM-BC) from the ANCOMBC R package (Lin and Peddada, 2020; Nearing *et al*., 2022). Linear discriminant analysis (LDA) effect size (LefSe) was performed to screen for differentially abundant predicted microbial function (MetaCyc pathways) across the four timepoints, based on an LDA threshold of 2.0 and a statistical significance cut-off of < 0.05 (All-against-all strategy; Segata *et al*., 2011).

Mantel test (Spearman correlation method) was employed to determine the correlation between the *I. nucleus* microbiome taxonomic structure and the oxidized PUFA products using their dissimilarity matrices across timepoints using the mantel() function in vegan package (Oksanen *et al*., 2022), which tested whether higher dissimilarity in microbiome was associated with higher dissimilarity in metabolite profiles. Since the oysters’ microbiome and metabolites were sampled from independent individuals, we averaged each microbial taxon and each metabolite concentration across replicates at each timepoints before generating a dissimilarity distance matrix between samples – Bray-Curtis (microbiome) and Euclidean (metabolites).

The GraphPad Prism software (San Diego, California, USA) was used to generate the figure for oyster’s body temperatures, individual oxidized PUFA products and the 10 dominant microbe’s relative abundance across the timepoints.

## Results

### Tidal dynamics and temporal changes in environmental and body temperatures

Oysters were regularly exposed to the air as a result of the tidal regime (Fig. 1A) and this temporal change was aligned to changes in the rock surface temperature across the day, with the highest temperature recorded at noon (45-50℃, Fig. 1A). The body temperature of the oysters varied temporally within a single emersion period (Fig. 1B; GLM *χ*^2^_(3)_ = 1073.9, p-value < 0.05), increasing from 32.11 ± 0.12 ℃ (mean ± SD) in the morning to 39.99 ± 0.95 ℃ (mean ± SD) at noon, after which body temperatures drop steadily to ∼ 35 °C in the afternoon. Besides early-afternoon (35.4℃ ± 0.46) and late-afternoon (35.18 ℃ ± 0.46), all pairwise comparisons showed significant differences in body temperatures between timepoints.

### Temporal assessment of oxidized PUFA products

Individual analyses of each oxidized PUFA products revealed that only four products (i.e., 5-F_2t_-IsoP, 10-F_4t_-NeuroP, 13-F_4t_-NeuroP, and 16-F_1t_-PhytoP) significantly varied between timepoints (Table 1, Fig. 2). These metabolites showed an increased in concentration at noon (10-F_4t_-NeuroP and 13-F_4t_-NeuroP) and post-noon (5-F_2t_-IsoP and 16-F_1t_-PhytoP) relative to morning, with varying degree in increment among these four metabolites. In particular, 5-F_2t_-IsoP and 16-F_1t_-PhytoP exhibited roughly 4-folds increase on average in concentration. There was a significant difference in the metabolite structural profile (PERMANOVA) (contributed by fifteen metabolites) between timepoints (*pseudo-F* = 2.17, p-value < 0.05; Fig. 3A), but PERMDISP indicated no significant difference in their dispersion (F-value = 0.69, p-value > 0.05; Fig. 3A). An additional PERMANOVA analysis with the four temporally-varied products showed a significant difference in the clusters between the morning metabolite profile and all the other timepoints during the emersion period (*pseudo-F* = 5.62, p-value < 0.05; Fig. 3B). Multivariate dispersion of the four oxidized PUFA products also differed between timepoints (F-value = 6.10, p-value < 0.05; Fig. 3B). With the exception of three pairwise comparisons (i.e., noon vs. early-afternoon, noon vs. late-afternoon and early-afternoon vs. late-afternoon), all the other timepoints showed significant differences in dispersion (p < 0.05). Combining both of these results, the metabolite structural profile at the morning timepoint had the smallest dispersion (smaller inter-interindividual difference) when only the four temporally-variable oxidized PUFA products were considered. However, when all products were included in the analysis, the overall group dispersion did not vary across the day.

**Fig. 2.**
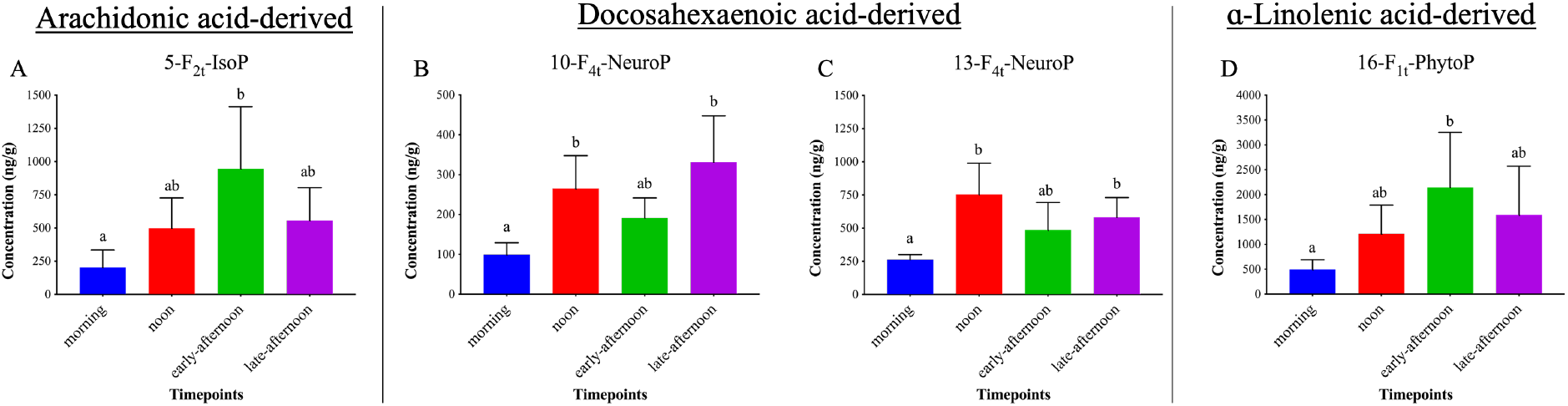
Four oxidized PUFA products exhibit variation in concentration across the emersion period. Bar graphs showing the four oxidized PUFA products deriving from their respective PUFA (labelled at the top of each panel, across different sampling timepoints in *I. nucleus*. IsoP: isoprostane; NeuroP: 19 europrostane; PhytoP: phytoprostanes. Data are presented as mean concentration and + 1 standard deviation (error bar) for each metabolite (n=5) at the given timepoints. Statistical significance (p-value < 0.05) is indicated if the letters above each bar are different.

**Fig. 3.**
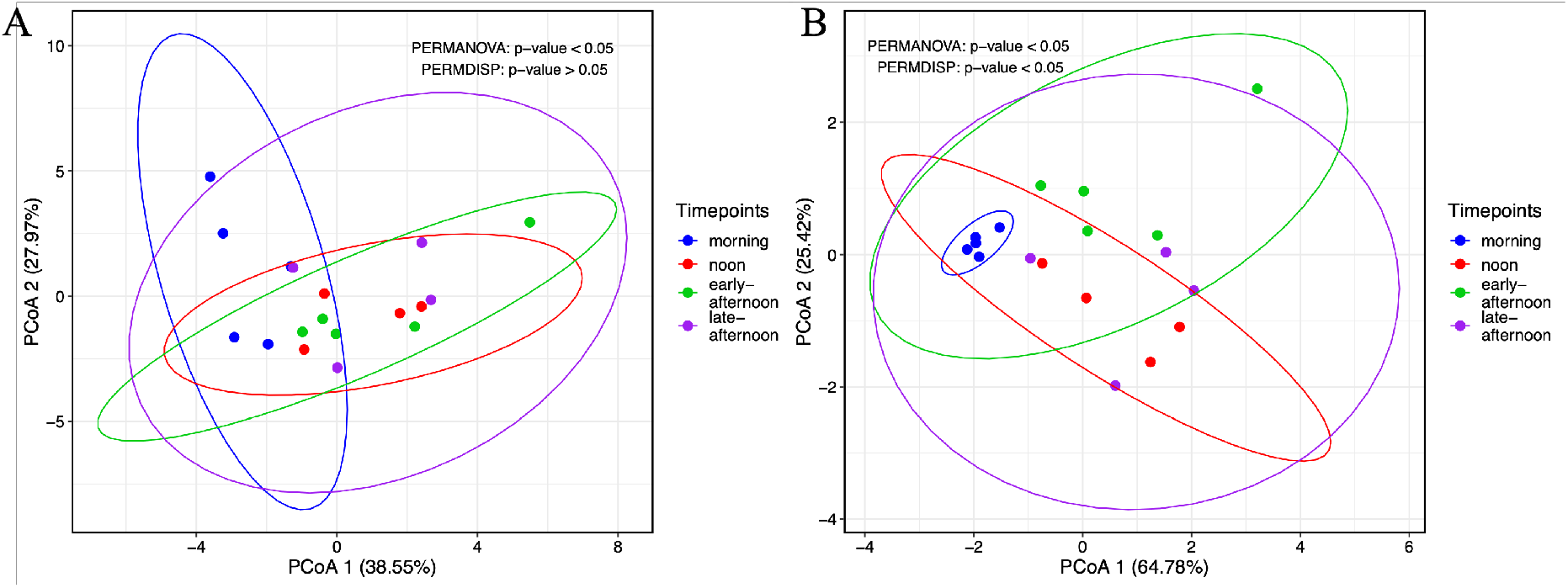
Oxidized PUFA products profile varies across the emersion period. PCoA based on Euclidean distance measured between samples from different timepoints based on (A) all oxidized PUFA products (B) four oxidized PUFA products: 5-F_2t_-IsoP, 10-F_4t_-NeuroP, 13-F_4t_-NeuroP, and 16-F_1t_-PhytoP. The colours of the datapoints and ellipses (95% confidence interval of the true multivariate means) indicate their corresponding timepoints.

**Table 1.** Statistical analysis result (ANOVA or GLM with gamma distribution) testing the effect of timepoints on the concentration of each of the oxidized PUFA products.

| PUFA | Metabolite | Test Statistic | p-value |
| --- | --- | --- | --- |
| Arachidonic acid | 5-F<sub>2t</sub>-IsoP | $F_{(3,14)} = 5.08$ | **0.04** |
| | 15-F<sub>2t</sub>-IsoP | $F_{(3,14)} = 1.72$ | 0.21 |
| Adrenic acid | 7-F<sub>2t</sub>-Dihomo-IsoP | $F_{(3,14)} = 3.10$ | 0.06 |
| | 17-F<sub>2t</sub>-Dihomo-IsoP | $GLM \chi^2_{(3)} = 0.66$ | 0.69 |
| Docosahexaenoic acid | 4-F<sub>4t</sub>-NeuroP | $F_{(3,14)} = 3.13$ | 0.06 |
| | 10-F<sub>4t</sub>-NeuroP | $F_{(3,14)} = 8.43$ | **< 0.01** |
| | 13-F<sub>4t</sub>-NeuroP | $GLM \chi^2_{(3)} = 27.46$ | **< 0.01** |
| | 20-F<sub>4t</sub>-NeuroP | $F_{(3,14)} = 1.23$ | 0.34 |
| α-Linolenic acid | 9-D<sub>1t</sub>-PhytoP | $F_{(3,14)} = 0.58$ | 0.64 |
| | 9-F<sub>1t</sub>-PhytoP | $GLM \chi^2_{(3)} = 7.92$ | 0.05 |
| | 16-F<sub>1t</sub>-PhytoP | $F_{(3,14)} = 3.72$ | **0.04** |
| | 16-B<sub>1t</sub>-PhytoP | $GLM \chi^2_{(3)} = 6.66$ | 0.08 |
| Eicosapentaenoic acid | 5-F<sub>3t</sub>-IsoP | $F_{(3,14)} = 2.49$ | 0.10 |
| | 15-F<sub>3t</sub>-IsoP | $GLM \chi^2_{(3)} = 0.32$ | 0.32 |
| | 8-F<sub>3t</sub>-IsoP | $F_{(3,14)} = 0.81$ | 0.51 |

### Temporal characterization of oyster-microbiomes

The 16S rRNA amplicon sequencing generated a total of 2,032,567 reads from 30 samples which were reduced to 1,126,884 after denoising and quality control. These reads were then used to characterize the microbial communities of *I. nucleus* across the different timepoints. Due to the low sequence counts, samples BRM3 (5612 reads) and DRM4 (7519 reads) were removed after each sample was rarefied to a sampling depth of 21,282 reads.

The ASV richness and Shannon-Wiener index of *I. nucleus* microbiome were significantly different between timepoints (ASV richness: _3,14)_ = 3.67; p-value < 0.05; Shannon-Wiener index: F_(3,14)_ = 3.76; p-value < 0.05; Fig. S2). However, no specific temporal trends were detected for all pairwise comparisons in ASV richness. Significant differences were, however, found for the Shannon-Wiener index between morning and noon timepoints. Oyster body temperature significantly influenced ASV richness and Shannon-Wiener indexes, showing an inverse relationship with microbiome alpha diversity (Fig. S3). Overall, morning timepoints showed the highest alpha diversity. This diversity then drastically declined at noon, with partial recovery at the last timepoint – late-afternoon.

The top 10 most abundant microbial taxa (at the family level) associated with *I. nucleus* in a descending order were *Mycoplasmataceae, Helicobacteraceae, Spirochaetaceae, Xenococcaceae, Rhodobacteceae, Rickettsiales* unknown*, Arcobactericeae, Caldilineaceae, Alphaproteobacteria* unknown and *Saprospiraceae* (Fig. 4). These top 10 families made up on average 57.09%, 80.88%, 81.42%, and 68.20% of the overall relative abundance in morning, noon, early-afternoon, and late-afternoon, respectively. Among them, only two families (*Helicobacteraceae* and *Arcobacteraceae*) showed significant differences in the relative abundance between timepoints based on the ANCOM-BC analysis (Fig. S4; *Helicobacteraceae*: morning vs early-afternoon; *Arcobacteraceae*: morning vs noon, and morning vs early-afternoon).

**Fig. 4.**
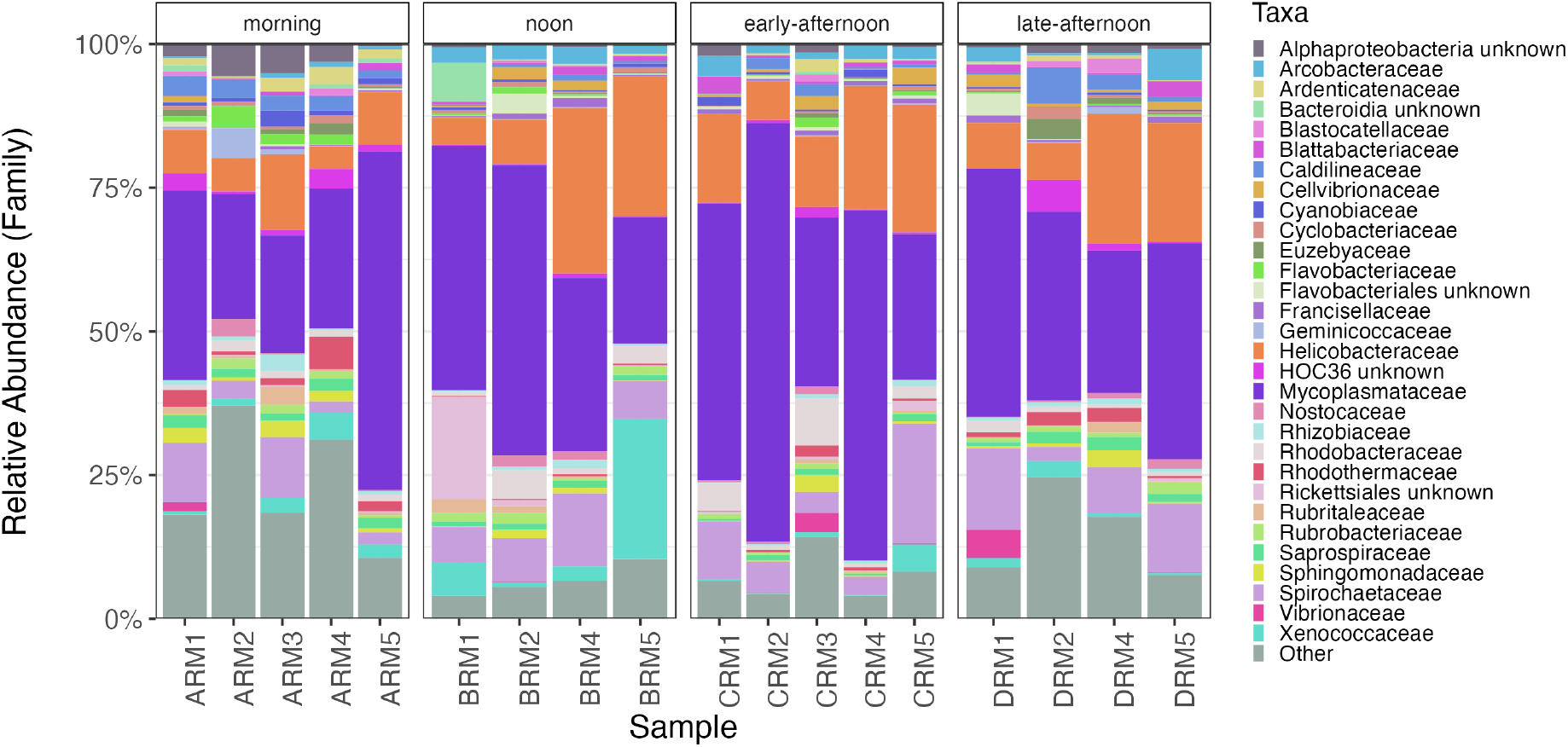
Relative abundance of top 30 most dominant microbial taxa associated with the *I. nucleus’s* gut microbiome at each sampling timepoints. ASV counts were aggregated into the family taxonomic level. Each bar represents a single individual replicate of oyster gut microbiome.

LefSe analysis identified 27 differentially abundant microbial functions (MetaCyc pathways; LDA threshold of 2.0) that characterises the microbiome at a particular timepoint (Fig. 5). The most distinctive difference in functions identified were between morning and late-afternoon timepoints, where the former showed a major enrichment in genes associated with biosynthetic pathway (and some biomolecules degradation pathways), while the latter had functions affiliated with degradation, utilization, and assimilation pathways, specifically in carbon metabolism.

**Fig. 5.**
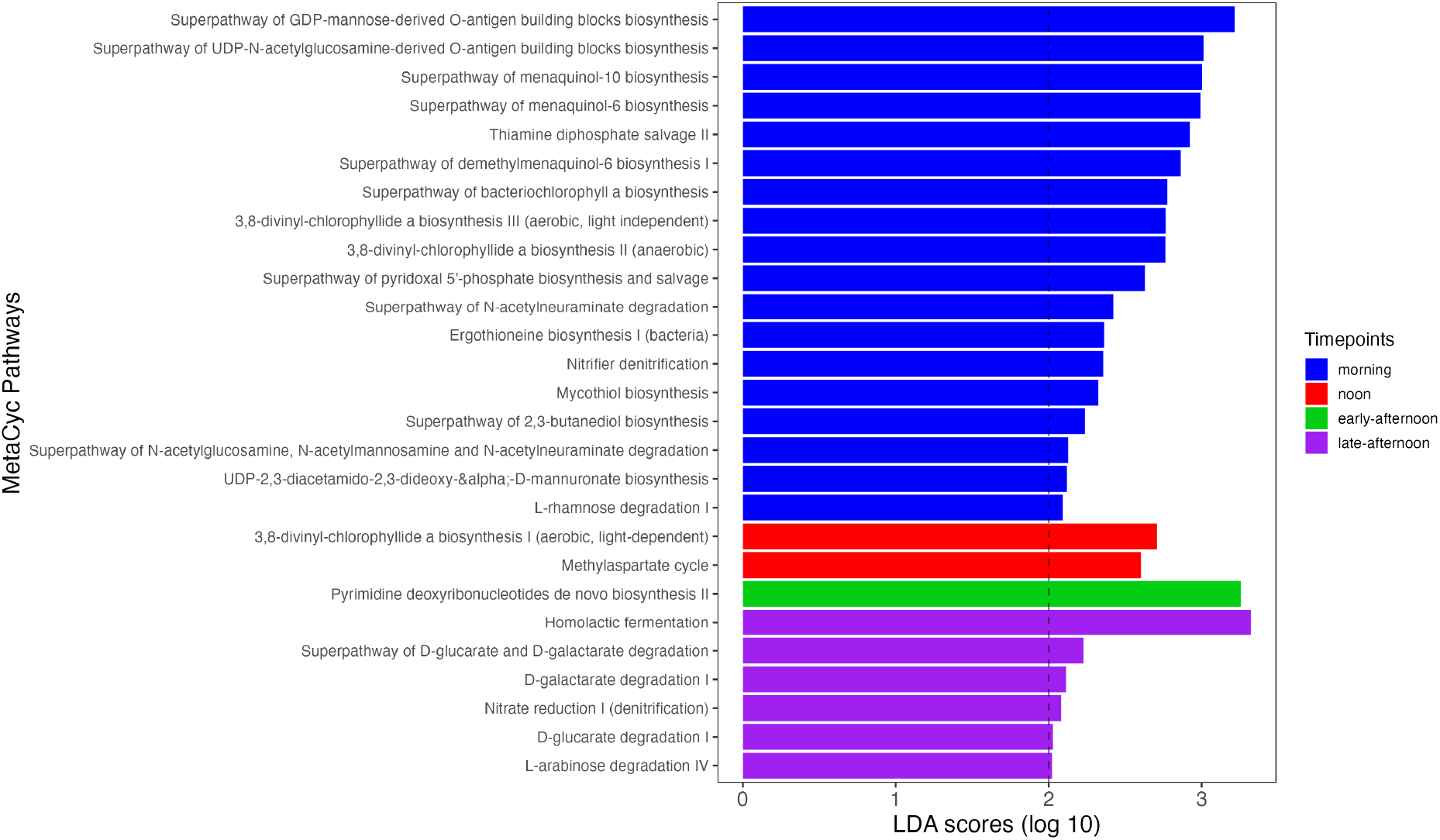
Differentially abundant microbial functions associated with a particular time point. LefSe analysis (LDA scores of > 2 and alpha of 0.05) showing 27 predicted functions (MetaCyc) pathways) that was identified as discriminative among the timepoints. Coloured bar represents functions that was derived from a particular timepoint. The LDA scores cut-off is depicted by the dotted lines.

Assessing the overall microbiome structure in taxonomy and functions with PERMANOVA analysis indicates no significant difference between timepoints (taxonomy: *pseudo-F =* 1.18, p-value > 0.05; function: *pseudo-F =* 1.49, p-value > 0.05;Fig. 6), but significant differences was detected in both of the microbial dispersion profiles (taxonomy: F-value = 3.55, p-value < 0.05; function: F-value *=* 3.74, p-value < 0.05; Fig. 6). Major differences in the taxonomic microbiome dispersion were between morning and early-afternoon, which corresponds to the pre-and post-noon period, respectively. For the functional microbiome, the pair early-afternoon-late-afternoon was the cause of the difference in dispersion. We found no Mantel correlation was observed between the microbiome taxonomic profile and the oxidized PUFA products profile, either with the 15 (all) or 4 metabolite groups (Table 2).

**Fig. 6.**
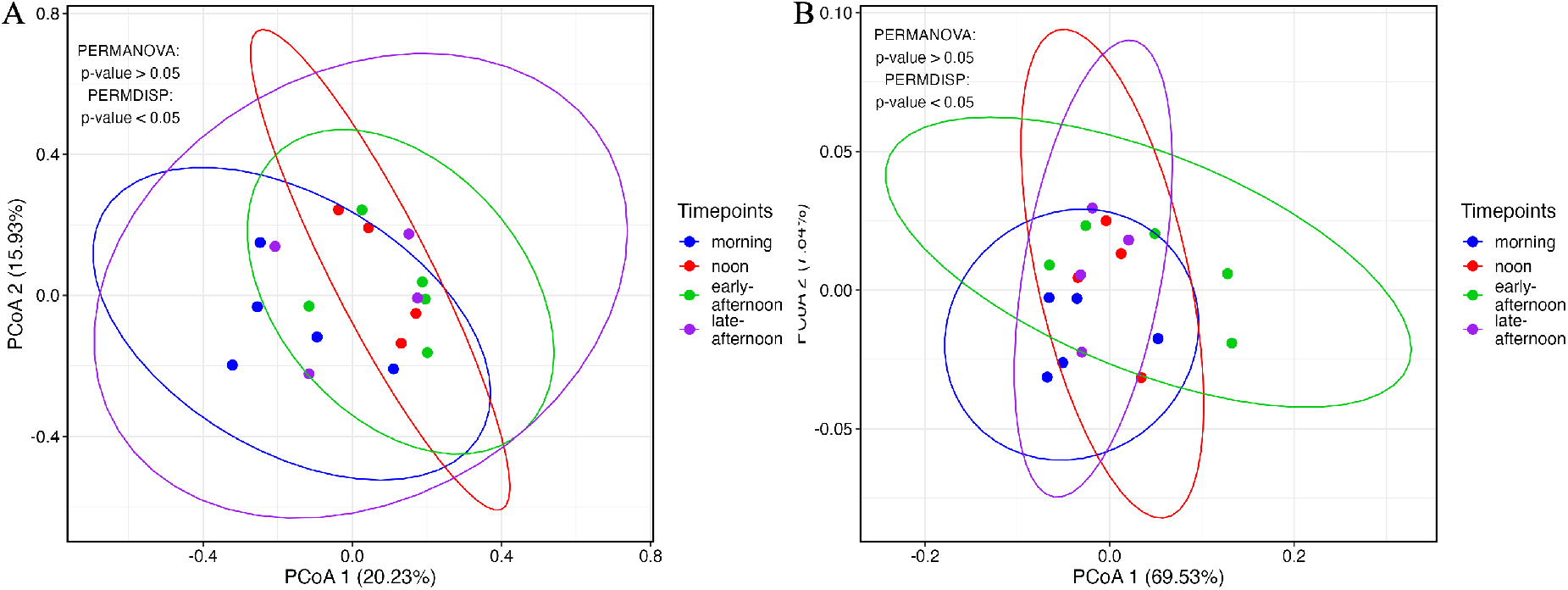
*I. nucleus*-associated microbiome taxonomic and functional structures exhibit variation in dispersion across the emersion period. PCoA based on the Bray-Curtis distance metric measured between samples from different timepoints, based on (A) taxonomy (ASV level) and (B) functions (MetaCyc pathways). The colours of the datapoints and ellipses (95% confidence interval of the true multivariate means) indicate their corresponding timepoints.

**Table 2.** Mantel’s spearman’s rank correlation I and p-value to investigate the association between the *I. nucleus* microbiome taxonomic structure (mean abundance of each taxon across all timepoints) and oxidized PUFA products profile (mean concentration across timepoints). The number of permutations performed was 23(maximum possible).

| Association with microbiome taxonomic structure | Mantel's r | p-value |
| --- | --- | --- |
| oxidized PUFA products (all) | 0.77 | 0.21 |
| oxidized PUFA products (Four)* | 0.54 | 0.25 |

\*The four metabolites were: 5-F<sub>2t</sub>-IsoP, 10-F<sub>4t</sub>-NeuroP, 13-F<sub>4t</sub>-NeuroP, and 16-F<sub>1t</sub>-PhytoP

## Discussion

Organisms living on the intertidal rocky shore regularly experience drastic changes in environmental temperature (Somero, 2002; Stillman, 2002; Harley, 2008). For many sessile ectotherms, this represents a physiological challenge due to their inability to escape from heat stress (e.g. seeking thermal refuge) and so, they rely on various intrinsic (e.g., metabolic and physiological plasticity) and extrinsic (e.g., host-associated microbiome; still underexplored) mechanisms to cope with temperature fluctuations (Lathlean *et al*., 2014; Giomi *et al*., 2016; Bang *et al*., 2018). As with other ectotherms, the body temperature of the high-supratidal oyster *Isognomon nucleus*, was highly influenced by changes in ambient temperature, emersion, and the reduction in oxygen availability and food during the low-tide period. Such changes particularly in temperature, are known to induce physiological stress and metabolic depression (Hui *et al*., 2020), with the associated oxidative stress (Hermes-Lima *et al*., 2015). Here, linked to these physiological conditions, we found adjustments in four major non-enzymatic oxidized PUFA products (i.e., 5-F_2t_-IsoP, 10-F_4t_-NeuroP, 13-F_4t_-NeuroP, and 16-F_1t_-PhytoP) that reflect temporal variation in the oxidative stress experienced by the oysters. When assessing the microbiome profile through time, there was a significant change in the alpha diversity, dominant microbial and specific functional enrichment. No clear temporal change in beta-diversity was detected in the gut microbiome, despite the significant variation in group dispersion. Our study found a lack of temporal correlation between the intrinsic (biochemical profile) and extrinsic (microbiome taxonomic structure) mechanisms studied. These findings suggest that environmental variation and/or host thermal condition (within a single emersion period), can drive independent physiological responses (biochemistry – oxidized PUFA products) and temporal dynamics in the gut microbiome of the tropical intertidal oyster *I. nucleus*. These independent responses may be important for the overall homeostasis of the oysters, and their ability to cope and survive in the thermally challenging tropical high-supratidal environment. However, manipulative experiments would be needed to confirm this hypothesis.

### The intrinsic mechanism underpinning oysters’ adaptive response to temporal environmental changes

For high-supratidal zone inhabiting ectotherms, metabolic depression is an adaptive response that allows them to cope with extreme high and stressful thermal conditions (Sokolova and Pörtner, 2001; Marshall *et al*., 2011). In a laboratory setting, Hui et al. (2020) demonstrated that the oyster *I. nucleus* consistently enters into metabolic depression (i.e., measured by heart rate) when body temperature approaches ∼ 37℃ (a value that is regularly surpassed on daily basis at around noon on tropical rocky shores). Based on this previous study, our data indicates that the oysters could have entered a state of metabolic depression when exposed during the low tide at noon time (*I. nucleus* noon temperature: ∼ 39.99 ± 0.95 ℃). After this, oysters resumed their metabolic activity when body temperatures decreased (i.e., late afternoon/evening).

Intertidal bivalves are known to also display intermittent gaping behaviour during periods of aerial exposure as a way to enhance oxygen intake and evaporative cooling, particularly when experiencing thermal stress (Lent, 1969; Mcmahon, 1988; Nicastro *et al*., 2010). In our study, all individuals shut their valves and no intermittent gaping behaviour was observed either during the high thermal stress or the recovering period (Hui, T.Y., unpublished data). Therefore, these oysters are expected to experience low oxygen availability within the shells, if not hypoxia. Under such condition, metabolic depression is a critical strategy for survival, reducing energy consumption and the elevation of ROS. This strategy is particularly relevant at noon where these oysters experienced the highest environmental temperature, which is usually associated with an increase in metabolic demand (Hermes-Lima *et al*., 2015; Jiang *et al*., 2020; Wang *et al*., 2022).

There were four metabolites (i.e., 5-F_2t_-IsoP (Isoprostane), 10-F_4t_-NeuroP (Neuroprostane), 13-F_4t_-NeuroP (Neuroprostane), and 16-F_1t_-PhytoP (Phytoprostane)) that showed visible temporal patterns, potentially associated with their metabolic regulation. Such patterns may be linked to the sensitivity of the metabolites to ROS changes in the oyster. The elevation of two metabolites at noon (10-F_4t_-NeuroP and 13-F_4t_-NeuroP) and two metabolites (5-F_2t_-IsoP, and 16-F_1t_-PhytoP) at post-noon timepoints, clearly indicates that oysters experienced a rise in the ROS level, a signal of oxidative stress (Hermes-Lima and Zenteno-Savín, 2002; Storey and Storey, 2004; Welker *et al*., 2013; Galano *et al*., 2017). The high levels of 16-F_1t_-PhytoP (Phytoprostane) metabolites, compared to other three oxidized PUFA products in the *I. nucleus,* imply that the PUFA ALA could be the primary target for lipid oxidative damage in this oyster. Our findings (i.e., changes in these metabolites over a short period of time ∼3 hours: morning – noon), support the observations by Williams and Somero (1996). These authors showed that over a short period of time (within hours), the gills of the intertidal *mussels Mytilus califonianus* (high intertidal zone) exhibit an alteration in the cellular membrane order/fluidity (i.e., phospholipid vesicles), in response to thermal variation encountered during a single period of tidal cycle. Thus, the rapid alteration in the biochemical profile observed in our study could be an important response for the survival of the oysters during periods of thermal stress and fluctuations in the intertidal rocky shore.

Further analysis with the overall structural profile with the four ROS-sensitive oxidized PUFA products showed a clear temporal variation and inter-individual variation in *I. nucleus*. The large inter-individual variation in the oxidized PUFA products profile at noon/afternoon relative to morning timepoint might be related to the thermal and oxidative stress conditions experienced by the oysters during those periods of elevated temperatures. This variability suggests inter-individual differences in physiological plasticity, regulation, and sensitivities to environmental stress. The result here also indicates that the polyunsaturated fatty acids (ARA, DHA, and ALA) from which these four oxidized PUFA products were derived, may play a key role as lipid metabolites that help to resolve inflammation and oxidative stress for the oysters to return to homeostasis (Roy *et al*., 2017; Lee *et al*., 2020). It is important to highlight here that the current understanding regarding the thermal-physiological roles of high PUFA ALA or the oxidized PUFA products in *I. nucleus* is still limited and that our interpretations are speculative and idiosyncratic to the specific animals and population studied. Moreover, our assessment was based on the observation and analysis that are known in clinical research and we remained carefully and strictly conservative in our discussion to just focus on the association between these products and the host oxidative stress level. Future studies would be dedicated toward elucidating the role of these molecules that may be essential to their host physiology and metabolism.

### The extrinsic mechanism underpinning the oysters’ response to temporal environmental changes

Host-associated microbiome diversity, structure, and functions are considered as important components of animal adaptation and acclimation to environmental change (Foster *et al*., 2017; Bang *et al*., 2018; Voolstra and Ziegler, 2020). Alterations in microbiome diversity can be associated with the direct influence of the host’s physiology and/or with host-independent environmental effects (Shiu *et al*., 2017; Li *et al*., 2018, 2022; Muñoz *et al*., 2019). In our study, the composition and diversity (ASV richness and Shannon diversity) of the oyster gut*-*associated microbiome varied with time which coincided with variation in the oyster’s body temperature, suggesting that the thermal condition experienced by the host may modulate the microbial alpha diversity via either direct/indirect physio-biochemical influences (Sorek *et al*., 2014; Pita *et al*., 2018; Liu *et al*., 2023). In the morning, the oyster’s gut had a significantly more diverse microbiome diversity, which may be linked to the less thermally stressed host (Hui *et al*., 2020), and/or to the filter-feeding activities before being exposed by the receding tide (Parris *et al*., 2019). However, based on the Shannon alpha diversity, at early post-noon condition, a major decline in diversity was observed, which may be associated with the thermal physiological stress experienced by the host (body temperature is within the metabolic depression range; Hui *et al*., 2020) and oxidative stress (indicated by the rise in the two oxidized PUFA products at noon and two at early post-noon condition). This decline in diversity is also a common pattern seen in many marine organisms under physiological stress (Green and Barnes, 2010; Li *et al*., 2018; Hartman *et al*., 2020). For example, Lokmer & Wegner (2015) showed that acute thermal stress and/or infection caused a marked decline in the alpha diversity of the microbiome in the oyster *Crassostrea gigas*. In another study by Li et al. (2018) observed an overall decrease in the gut microbial alpha diversity associated with the mussel, *Mytilus coruscus*, after exposing to higher temperature. Such observation offer support to our result where the relatively higher temperature at noon may have driven the decline in the oyster-associated microbial diversity.

Whilst the temporal changes exhibited by the *I. nucleus* microbiome and their association with changes in the oysters’ body temperatures suggest that the microbiome composition may be actively regulated by the host, differential sensitivities of certain bacterial taxa to external environmental changes may also be an important mechanism driving the temporal variation in the microbiome composition. *Arcobacteraceae* and *Helicobacteraceae* are the only two dominant family-level microbial taxa associated with the temporal thermal/oxidative stress variation of the host. In particular, there was an elevated relative abundance of these two bacteria groups at noon and/or at the afternoon timepoints, which may suggest that their abundances during these periods were promoted by specific host/environmental conditions, facilitating their growth probably by reducing competition (Garren *et al*., 2016; Shaver *et al*., 2017; Sharp and Foster, 2022). *Arcobacteraceae* is a group of common opportunistic bacteria found in marine bivalves and is known to be associated with disease-causing bacteria such as *Vibrio* (Lokmer and Wegner, 2015; de Lorgeril *et al*., 2018). In many studies, the elevation of the *Arcobacteraceae* has been associated with the thermal stress condition experienced by the host (Lokmer and Wegner, 2015; Shiu *et al*., 2017; Neu *et al*., 2021). Members of *Arcobacteraceae* are also known to be microaerophilic as they tend to grow better under a low-oxygen environment (Vandamme and De Ley, 1991). This may explain the dominant presence of *Arcobacteraceae* within the oysters’ microbiome due to the prevailing hypoxic and thermally stressful conditions as experienced by the oyster hosts during emersion. Similarly, the presence and abundance *Helicobacteraceae* can be associated to oxygen levels in the marine environment. For instance, certain members of this family are capable of oxidising sulfide, and thus, they might contribute with a detoxification function to the oyster host (Kodama and Watanabe, 2004; Nakagawa *et al*., 2017; Neu *et al*., 2021). The accumulation of sulfide can be detrimental or lethal to marine bivalves (Le Moullac *et al*., 2008), and it is typically associated with hypoxic stress (Le Moullac *et al*., 2008; Pal *et al*., 2018). Such stress is hypothesised to occur in *I. nucleus* during aerial exposure, as no intermittent gaping behaviour was detected. When oxygen is low, sulfide production is promoted by the activity of hydrogen sulfide-producing bacteria such as *Desulfobacterota* members (Flannigan *et al*., 2011; present in the I. nucleus; Dordević *et al*., 2021; Pimentel *et al*., 2021). It is unclear, however, the extent to which the dynamics in the abundance of sulfide-producing bacteria and *Helicobacteraceae* members (with potential detoxification role), are functionally interconnected. While this seems likely, we are unaware of any study exploring this functional link, and it would be important to investigate such species’ interaction and its effects on the physiology and fitness of the host.

Variation in the microbiome functional activity is commonly associated with the functional changes occurring within the host (e.g., dietary changes; Mekuchi *et al*., 2018; Masasa *et al*., 2023) and/or in the external environment (e.g., temperature; Brothers *et al*., 2018; Li *et al*., 2022). Our study is in-line with this, in showing that there were several enriched functions that vary through time, potentially attributed to either one or both of those two factors. Specifically, in the morning timepoints, we detected a major proportion of predicted microbiome function in biosynthetic pathways associated with carbohydrate, cofactor, carrier, vitamin and secondary metabolites biosynthesis. This suggested that, in the morning, the host gut had a considerable dietary resources for the building of these organic/inorganic molecules in microbes to support cellular maintenance and growth and/or actively contributing to host nutritional metabolism (David *et al*., 2014; Maurice *et al*., 2015; Dubé *et al*., 2019). On contrary, post-noon conditions were significantly dominated by compound/molecular degradation pathways associated with carbon metabolism. This makes sense since in the post-noon conditions (post-heat stress), the host are likely in a “recovery” that could incur a high energetic requirement for various cellular and physiological repair and maintenance purposes (Storey and Storey, 2004; Sokolova *et al*., 2011). The breakdown of these carbon compounds by the microbes can provide as an energy source to meeting the metabolic demand in the host and microbes (Flint *et al*., 2012; Tremaroli and Bäckhed, 2012; Den Besten *et al*., 2013). However, it is unclear if these predicted microbial functional enrichments are directly contributing to the host dietary metabolism (e..g., nutrient absorption; English *et al*., 2023) or simply fulfilment of need for the microbes themselves (host-independent) to cope and survive to the prevailing conditions (i.e., host and/or external environmental influence).

Host tolerance and resilience to environmental stress are known to be conditioned by the stability/flexibility of its associated microbiome. While we found no major differences in microbiome beta diversity throughout the emersion period (based on PERMANOVA), there was a significant difference in dispersion between the timepoints. This suggests that the temporal change in the environment and host could influence the inter-individual variation in their microbial profile. Such temporal differences (particularly in alpha diversity) could potentially be explained by the ability of the host to control/regulate the microbiome during a disturbance event (Sharp *et al*., 2017; Pita *et al*., 2018; Rocca *et al*., 2019; Rädecker *et al*., 2022). Alternatively, temporal changes in the microbiome may reflect the result of its own control in response to changing and stressful environments, rather than the host control/regulation (Soen, 2014). These findings contrast with other studies showing that microbiome stability is essential for host tolerance and acclimation to environmental challenges such as thermal stress. For example, Santoro et al. (2021) demonstrated that the stability of inoculated “beneficial microorganisms” in the coral *Mussismilia hispida* provided a thermal protection effect, reducing heat stress and preventing the associated mortality (40% increase in the survival rate) of the hosts. Regardless whether it is host control or an independent reultation of the microbiome, the resilience of this extrinsic mechanism may contribute to the survival of *I. nucleus* in the thermally extreme tropical intertidal environment (Bang *et al*., 2018; Aktipis and Beltran, 2021).

### No correlation between microbiome structure and oxidized PUFA products profile

Over the decades, many studies have documented the role and action of PUFA (e.g., as probiotics) in modulating dynamics in the gut microbiome of different animals (Huyben *et al*., 2020; Fu *et al*., 2021; Lau *et al*., 2022; Ma *et al*., 2022). Hence, it is expected that stress-driven changes in these biochemicals and their by-products would influence the oyster gut microbiome. However, we found no correlation between the oxidized PUFA products (averaged across all tissues) and the gut microbiome taxonomic structure in *I. nucleus*, despite the temporal and inter-individual variation in the oxidized PUFA products profile across the emersion period. The lack of association between the microbial and oxidized PUFA products profiles of *I. nucleus*, suggests two possibilities that are not mutually exclusive. Firstly, as mentioned previously, physiological/biochemical and microbiome profiles may have different sensitivities to changes in environmental conditions. Secondly, such microbiome profiles and biochemical responses may not show strong variation and/or alignment because of local adaptation. Here, the thermal window and oxygen levels experienced by the oysters, although extreme for many ectotherms, can be considered within the tolerance range of the species. Several studies have shown that the microbiome-metabolites correlation is detectable only under severe post-disturbance events and unhealthy host conditions (Nguyen *et al*., 2021; Pimentel *et al*., 2021; Walke *et al*., 2021; Mallott *et al*., 2022), which go beyond tolerance levels. Thus, in this study, oysters may be in a condition where stress is within the tolerance range, and thus no correlation can be observed between these two components (i.e., host biochemical profile and microbiome structure). Future studies aiming to explore the functional regulation and interplay between intrinsic and extrinsic mechanisms in ectotherms living in the thermally extreme and fluctuating environment like in this study, the intertidal rocky shore, require the assessment of both components when animals are closer to, or surpass their physiological tolerance limits at sublethal levels.

### Study limitations

While our findings are noteworthy, here we discuss some study limitations and potential recommendations to address them. First, temperature-modulated dynamics in host-associated microbial communities and biochemical profiles across the day lacked temporal replication. Although the selected timepoints are eco-physiologically relevant for the oysters, the consistency and repeatability of the trends observed can only be assessed by sampling at different and independent days (i.e., two or more non-consecutive days). Second, the timepoints selected, although extreme for most ectotherms on Earth, fell within a thermal window that represents acute low physiological stress to the oysters. Major and divergent differences in extrinsic and intrinsic responses might potentially be detected under acute high or chronic physiological stress when body temperatures reach the maximum temperature documented for this species 50-52°C in the field (Fig. 1). Third, the existence of an interacting drivers that may influence the microbiome altogether, such as, temperature, hypoxic condition (no gaping in the oyster) and the emersion duration. This makes our study difficult to distinguish the influence of a sole driver in the field. Fourth, the oyster microbiome can exhibit tissue-specific profiles associated to functional differences (Dubé *et al*., 2019). Here, our focus was the gut microbiome as its captures a large diverse microbial sets associated with their host (King *et al*., 2012; Pierce and Ward, 2018; Dubé *et al*., 2019), and several documented findings showing their strong and direct signal of their host physiological states (Li *et al*., 2018; Pierce and Evan, 2019; Fassarella *et al*., 2021; Stevick *et al*., 2021). However, other tissues such as the haemolymph or the gills of the oysters can provide complementary information regarding the interaction between microbial communities and their host under environmental stress (Lokmer and Wegner, 2015; Scanes *et al*., 2021). Despite these limitations, our study provides for the first-time empirical evidence of the intrinsic and extrinsic mechanisms underpinning the temperature tolerances and ecological success of tropical high-supratidal zone inhabiting ectotherms. Future studies can integrate our recommendations to provide more robust and independent assessments of temporal dynamics in temperature-modulated functional responses of ectotherms in extreme environments.

## Conclusion

Intrinsic (i.e., non-enzymatic oxidized PUFA products) and extrinsic (i.e., microbiome) mechanisms linked to the ecophysiology and tolerance/sensitivity of the oyster *Isognomon nucleus*, showed temporal variation, indicative of thermal and oxidative stress. Such trend showed a temporal association with the environmental and body temperatures during the tidal cycle where oysters were fully emersed. We observed a dynamic pattern in the oyster’s gut microbiome in terms of the alpha diversity, specific individual microbial taxa and functions that varied along this temporal and environmental cycle. However, limited changes were detected in the overall structure and function of the associated microbial communities. When assessing both the intrinsic and extrinsic mechanisms together, we found no clear correlation between the oxidized PUFA products profile and the microbiome profile, suggesting an independent influence from the environment and/or the host thermal condition on these two profiles. To our best knowledge, this is the first study to report the use of non-enzymatic oxidized PUFA products as a proxy for oxidative stress levels in intertidal animals and profiling short-term dynamics of the host-associated microbiome in an intertidal environment. Future investigations should aim to explore the interplay between these biomolecules and microbial patterns when stressful environmental conditions are unpredictable and beyond tolerance windows of the hosts. Moreover, it is important to understand whether these functional links between intrinsic and extrinsic mechanisms are modulated by local adaptation and can be shared among other intertidal organisms.

## Author Contributions

JDGE and BSA conceived the idea and designed the work. BSA carried out most of the work: data collection, sample processing, statistical analyses, and data presentation. AWTC, HTY GAW, and MG were involved in sample and environmental data collection. LKS performed the oxidized PUFA products extraction and processing. TD supplied the in-house synthesised oxidized lipids standards for PUFA metabolites quantification. JTY and TD were involved in analysing the metabolite data. BSA led the manuscript writing with continuous feedback from JDGE. All authors provided feedback on the final version of this work, and accepted responsibility for the entire content of this manuscript.

## Acknowledgement

We would like to thank The Swire Group of Companies, especially Ms. Tina Chan and Cathay Pacific staff for arranging air transportation of equipment to and from Thailand from Hong Kong

## Data availability

All raw data, and scripts will be made available once published.

## Supplementary information

**Fig. S1.**
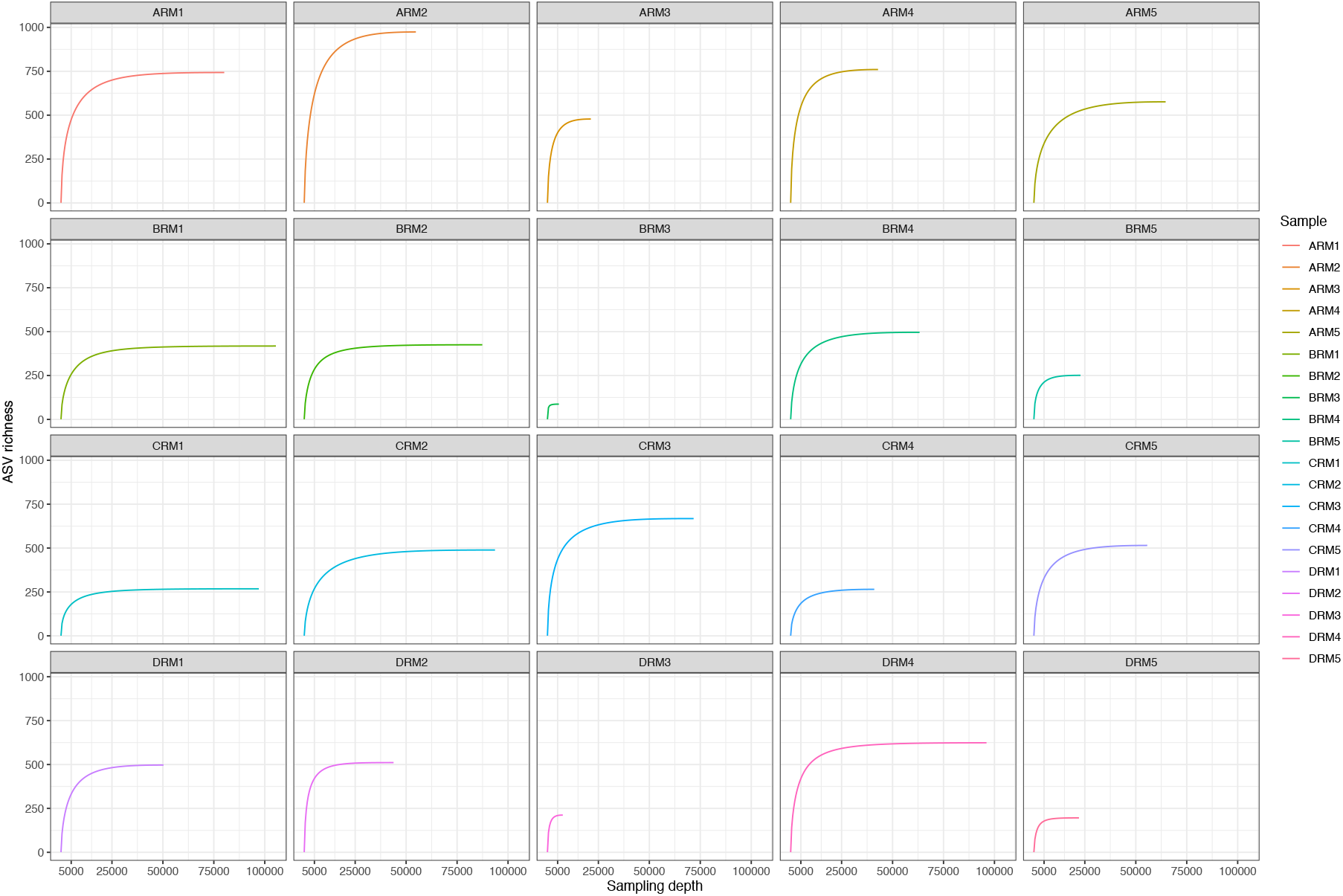
Rarefaction curves illustrating the ASVs captured against the sampling depth which was generated for each sample using their corresponding quality-filtered V3-V4 16SrRNA sequences.

**Fig. S2.**
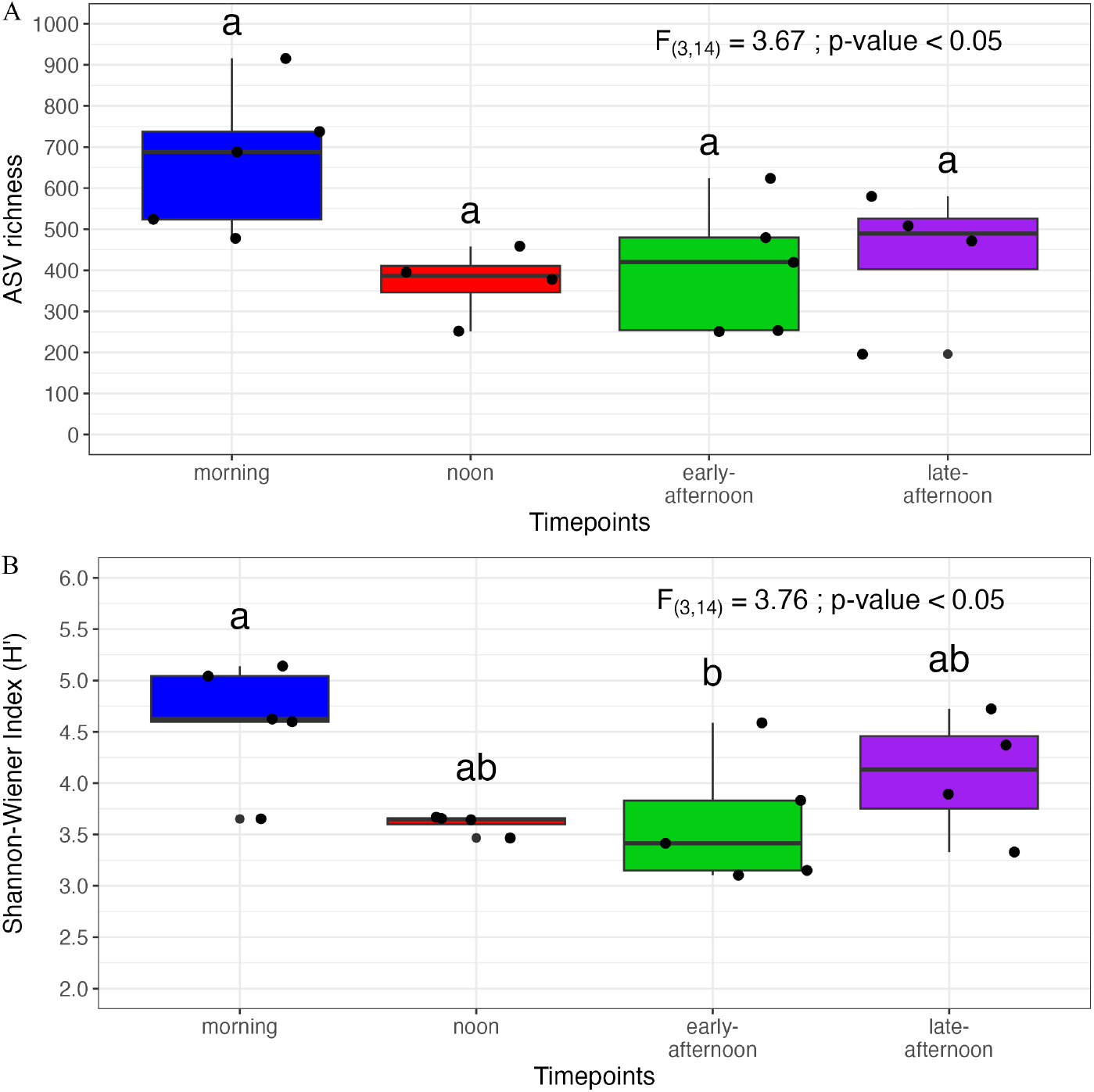
*I. nucleus-*associated microbiome ASV richness and Shannon-Wiener diversity across the emersion period. Boxplot showing the alpha diversity index of (A) ASV richness, and (B) Shannon-Wiener index (H’) of the oyster*-*associated microbiome at different timepoints. (Centre line represents the median; box limits indicate the first and third quartiles; the lower and upper whiskers indicate the smallest and largest values that within 1.5 times of the IQR (Inter-quartile range) from the first and third quartiles, respectively. The data points shown in each timepoint is replicate values. Statistical significance (p < 0.05) is indicated if the letters above each boxplot are different between any two timepoints.

**Fig. S3.**
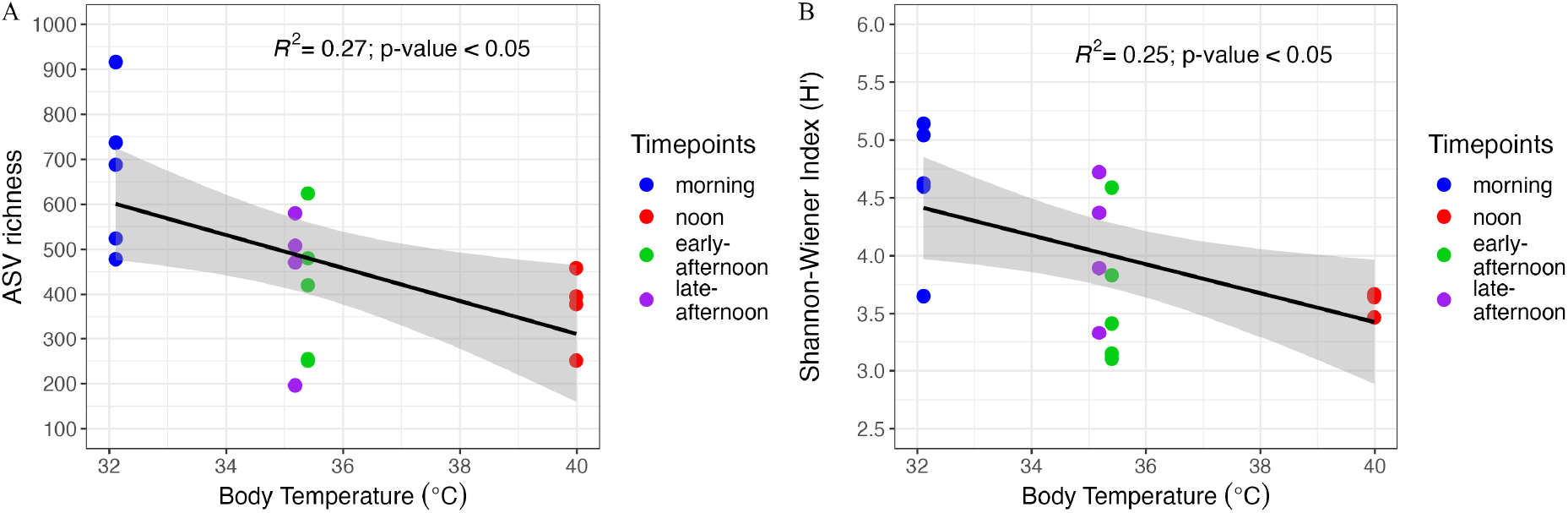
Variation in the alpha diversity indices of the *I. nucleus-*associated microbiome can be explained by the host body temperature. Linear regression graph showing the relationship between average body temperature (from 10 random independent replicates collected at each timepoint) and (A) ASV richness (B) Shannon-Wiener index (H’). The datapoints shown in each panel were the samples’ replicates (n= 4-5) from their corresponding timepoint. The shaded grey area indicates a 95% confidence interval for the predicted regression line. Statistical result (least square regression analysis) is shown in the upper right within each panel.

**Fig. S4.**
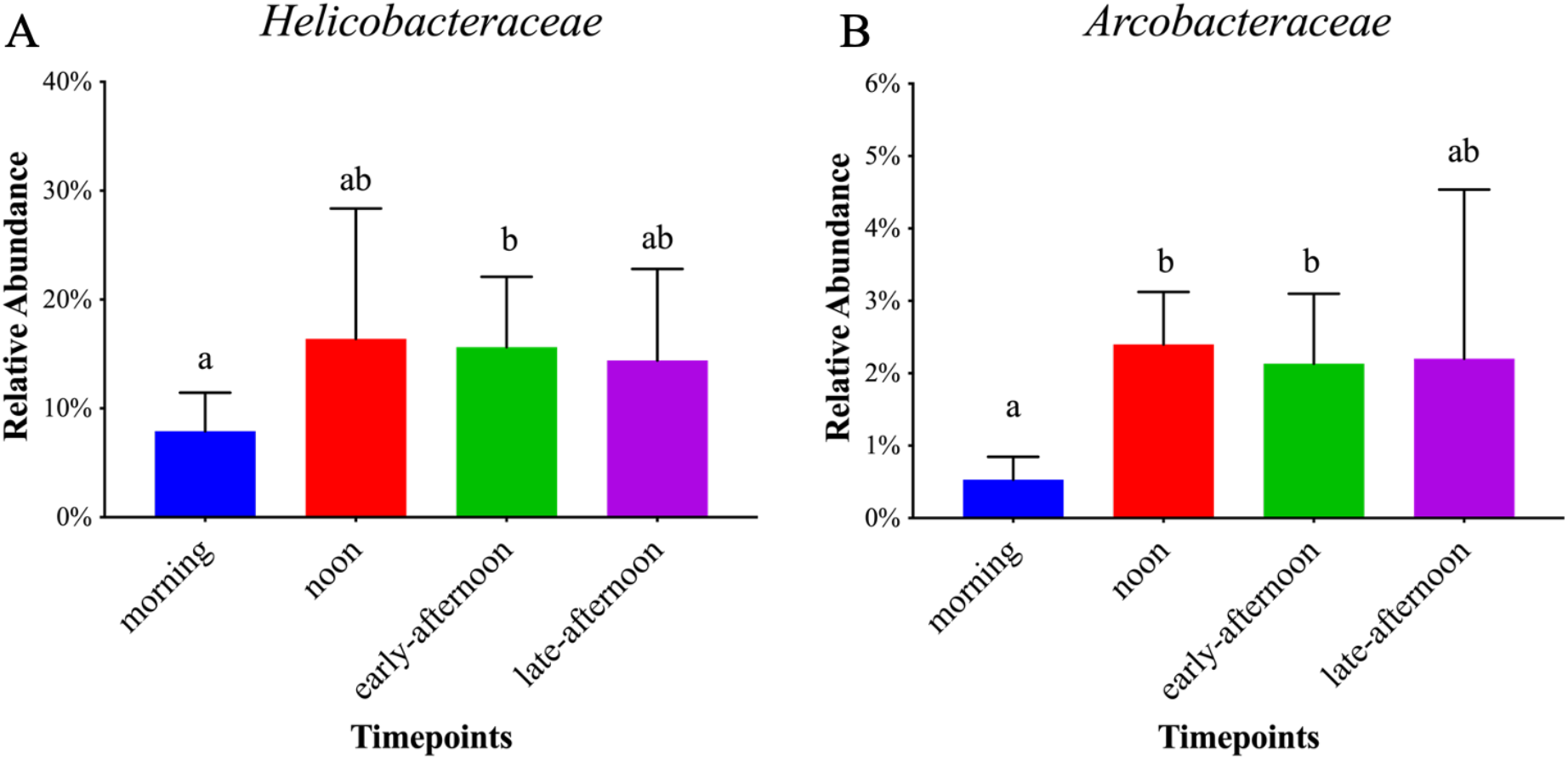
*Helicobacteraceae* and *Arcobacteraceae* mean relative abundance vary across the emersion period. Bar graphs showing the two dominant family level taxa in *I. nucleus*’s gut microbiome at each timepoint: (A) *Helicobacteraceae* (B) *Arcobacteraceae*. Data are presented as mean relative abundance and + 1 standard deviation (error bar) for each taxon at the given timepoints. Statistical significance (p-value < 0.05) is indicated if the letters above each bar are different.

**Table S1.** Metadata describing the *Isognomon nucleus* samples’ collected information.

| Date | Time | Timepoints label | Replicates | Sample name | Body (mm) |
| --- | --- | --- | --- | --- | --- |
| 22.05.2019 | 0915 | morning | 1 | ARM1 | 12.9 |
|  |  |  | 2 | ARM2 | 13.5 |
|  |  |  | 3 | ARM3 | 15.2 |
|  |  |  | 4 | ARM4 | 12.5 |
|  |  |  | 5 | ARM5 | 13.7 |
|  | 1230 | noon | 1 | BRM1 | 11.7 |
|  |  |  | 2 | BRM2 | 11.3 |
|  |  |  | 4 | BRM4 | 11.9 |
|  |  |  | 5 | BRM5 | 12.4 |
|  | 1612 | early-afternoon | 1 | CRM1 | 11.7 |
|  |  |  | 2 | CRM2 | 12.9 |
|  |  |  | 3 | CRM3 | 13.0 |
|  |  |  | 4 | CRM4 | 12.0 |
|  |  |  | 5 | CRM5 | 11.7 |
|  | 1740 | late-afternoon | 1 | DRM1 | 11.8 |
|  |  |  | 2 | DRM2 | 10.9 |
|  |  |  | 4 | DRM4 | 11.1 |
|  |  |  | 5 | DRM5 | 11.9 |

**Table S2.** Description of oxidized poly-unsaturated fatty acids (PUFA) products PUFA Oxidized PUFA products.

| PUFA | Oxidized PUFA products |
| --- | --- |
| <u>A</u>rachidonic <u>a</u>cid (ARA) | 5-F<sub>2t</sub>-IsoP |
|  | 15-F<sub>2t</sub>-IsoP |
| <u>A</u>drenic <u>a</u>cid (AdA) | 7-F<sub>2t</sub>-Dihomo-IsoP |
|  | 17-F<sub>2t</sub>-Dihomo-IsoP |
| <u>D</u>ocosa<u>h</u>exaenoic <u>a</u>cid (DHA) | 4-F<sub>4t</sub>-NeuroP |
|  | 10-F<sub>4t</sub>-NeuroP |
|  | 13-F<sub>4t</sub>-NeuroP |
|  | 20-F<sub>4t</sub>-NeuroP |
| <u>α</u>-<u>L</u>inolenic <u>a</u>cid (ALA) | 9-D<sub>1t</sub>-PhytoP |
|  | 9-F<sub>1t</sub>-PhytoP |
|  | 16-F<sub>1t</sub>-PhytoP |
|  | 16-B<sub>1t</sub>-PhytoP |
| <u>E</u>icosapentaenoic <u>a</u>cid (EPA) | 5-F<sub>3t</sub>-IsoP |
|  | 15-F<sub>3t</sub>-IsoP |
|  | 8-F<sub>3t</sub>-IsoP |

